# Api5-FGF2 regulates the transformation of breast epithelial cells via PDK1/Akt and Ras/MAPK/ERK signalling

**DOI:** 10.64898/2026.02.17.706358

**Authors:** Anisha Goyal, Mayurika Lahiri

## Abstract

The equilibrium between cell death and cell division is crucial for maintaining tissue homeostasis in a multicellular organism. Apoptosis plays an essential role in preserving homeostasis and hence occurs in a coordinated manner. However, inhibition of apoptosis is one of the hallmarks of cancer. Apoptosis Inhibitor 5 (Api5), an anti-apoptotic protein, is upregulated in various cancers, including ovarian, bladder, cervical, and lung cancers. Studies have demonstrated that altered expression of Api5 leads to the transformation of non-tumorigenic breast epithelial cells. However, the mechanism regulating this process is not well-elucidated. Our study demonstrates that overexpression of Api5 increased FGF2 (Fibroblast Growth Factor 2) levels both at protein and transcript levels. We studied the mechanistic details of changes in morphology, proliferation, and polarity observed upon FGF2/FGFR1 deregulation in Api5-overexpressing cells. Deciphering the signalling mechanism underlying Api5-FGF2-mediated breast tumorigenesis revealed that the PDK1/Akt and Ras/MAPK/ERK pathways regulated multiple transformation phenotypes. PDK1/Akt enhanced proliferation and altered morphology during initial stages, whereas Ras/MAPK/ERK regulated polarity disruption, proliferation, and reduced apoptosis during later stages of morphogenesis. In conclusion, this study provides insights into the signalling mechanism regulating the transformation phenotypes associated with Api5 overexpression in a non-tumorigenic breast epithelial cell line.

## Introduction

According to GLOBOCAN 2020, breast cancer is one of the leading causes of death in females all over the world [1]. In 2011, Hanahan and Weinberg proposed ten hallmarks of cancer, which are the characteristics a cell acquires during its transition from a normal to a transformed state. One of the hallmarks is resistance to cell death, which is acquired by inhibiting apoptosis [2]. Apoptosis or programmed cell death is a primary mechanism regulated by various pro-apoptotic and anti-apoptotic proteins. Pro-apoptotic proteins initiate apoptosis, whereas anti-apoptotic proteins inhibit it. The balance between the two determines whether a cell will undergo apoptosis [3].

Api5 or apoptosis inhibitor 5, has been reported as a plausible oncogene. It is also known as AAC-11 (Anti-apoptotic clone 11) [4], FIF (FGF2 interacting factor) [5], and MIG 8 (Migration Inducing Gene 8), and is a nuclear-localised, 55 kDa protein [6]. The N-terminal of Api5 consists of an LxxLL motif, whereas the C-terminal consists of LZD (Leucine Zipper Domain) and NLS (Nuclear Localisation Signal) [6] domains. Api5 was discovered as a cDNA clone that promoted cell survival upon serum starvation [4]. Over the years, various functions of Api5 have been reported. It was shown to regulate the transcription of E2F1-mediated genes, thereby inducing apoptosis and cell division [7] [8]. Api5 also regulates apoptosis by inhibiting caspase 2 homo dimerisation as well as FGF2/FGFR1-mediated Bim (a pro-apoptotic protein) degradation [9] [10].

FGF2 (Fibroblast Growth Factor 2) was initially isolated from the brain and pituitary gland as a growth factor for fibroblasts [11]. It exists in 5 isoforms, categorised into two subclasses: HMW FGF2 (High molecular weight FGF2), which comprises 22, 22.5, 24, and 34 kDa isoforms, and LMW FGF2 (Low molecular weight FGF2), which comprises the 18 kDa isoform [12]. Its primary mechanism of action involves translocation into the cytosol and binding to the transmembrane FGFR1 (Fibroblast Growth Factor Receptor 1). This leads to autophosphorylation of the receptor’s tyrosine kinase domain, thereby recruiting the adaptor protein FRS2α (FGF Receptor substrate 2). FRS2α, once recruited, gets phosphorylated and further activates downstream signalling [13].

Api5 has been shown to physically interact with LMW FGF2, which is essential for mRNA export, specifically those that respond to eIF4E, such as cyclin D1 and cMyc [14]. Api5 and FGF2 were also found to be upregulated in various cancers, including cervical cancer [15], B-cell chronic lymphoid leukaemia [16], and breast cancer [17]. Moreover, Api5 was also found to regulate cancer stem cell-like properties of cervical cancer cell line via FGF2-mediated NANOG axis [15].

Previous studies from our lab have reported that overexpression of Api5 leads to the transformation of breast epithelial cells, affecting proliferation, apico-basal polarity, apoptosis, and sphericity, among other parameters [17]. However, the molecular mechanism regulating this was not well elucidated. Therefore, this study investigated the signalling pathway regulating Api5 overexpression-mediated transformation using the MCF10A isogenic series as a model system [18] [19], consisting of MCF10A (a non-tumorigenic breast epithelial cell line), MCF10AT (a premalignant cell line), and MCF10CA1a (a malignant cell line). We report that overexpression of Api5 in MCF10A activates FGF2/FGFR1 signalling, leading to activation of the PDK1/Akt pathway during the initial stages of morphogenesis, thereby regulating altered morphology and enhanced proliferation. This signalling later switches to the FGF2-mediated Ras/MAPK/ERK pathway during later stages of morphogenesis to regulate proliferation, disrupted polarity, altered morphology and lumen filling. Our findings provide insights into Api5-FGF2-mediated transformation of breast epithelial cells, which can be further used to develop therapeutic approaches.

## Results

### Api5 positively correlates with FGF2 in breast cancer

To investigate the correlation of API5 and FGF2 in breast cancer, *in silico* analysis was performed using the TCGA (The Cancer Genome Atlas) database. The plot in Figure 1a, demonstrates a positive correlation between API5 and FGF2 at the transcript level in breast cancers. When analysed across the different stages of breast cancer, only Stage II showed a positive correlation between the two proteins (Figure 1b). When similar analyses were performed across breast cancer molecular subtypes, only Luminal B showed a positive correlation between API5 and FGF2 (Figure 1c).

**Figure 1:**
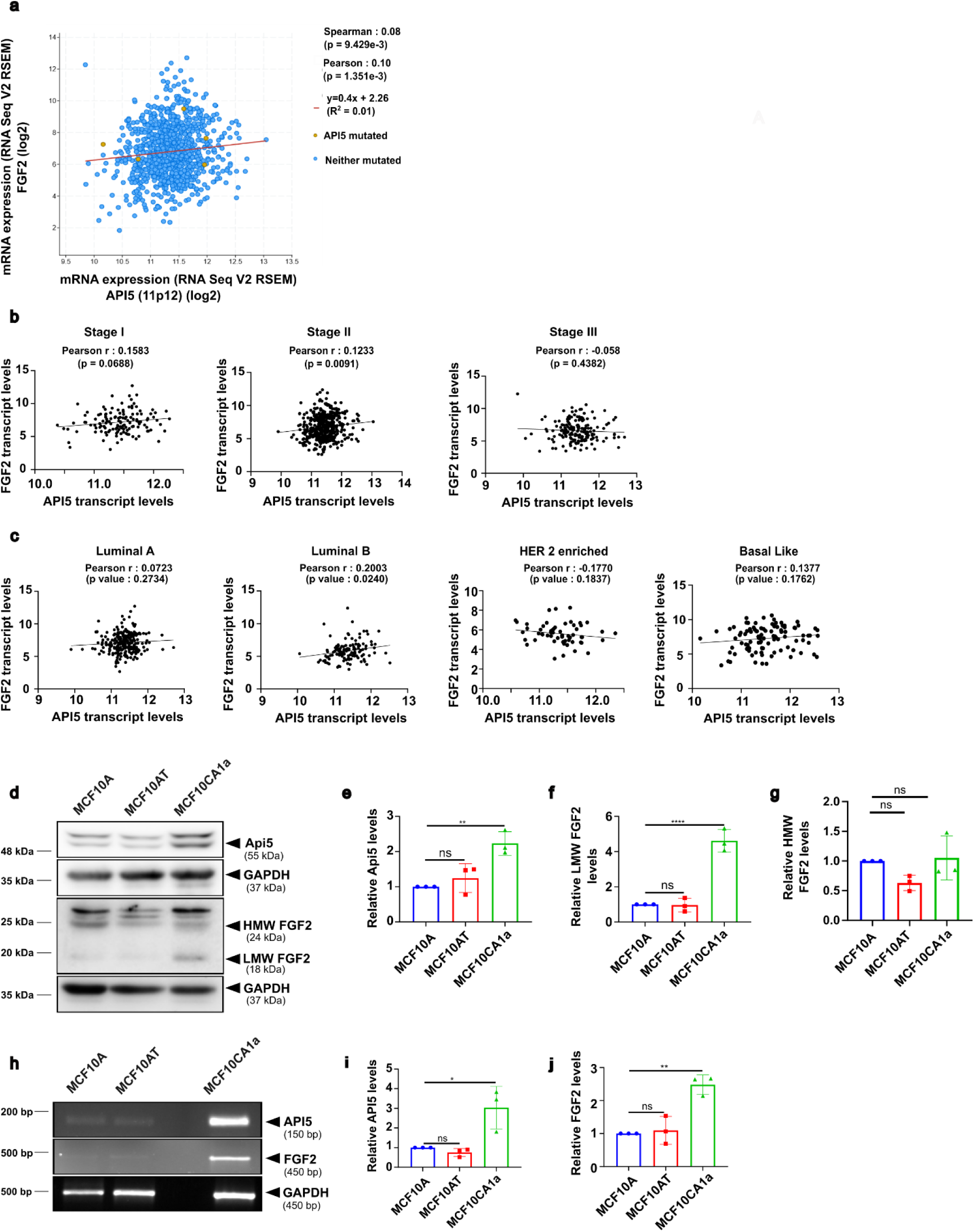
Api5 and FGF2 positively correlates in breast cancer: (a) mRNA levels of API5 and FGF2 shows positive spearman correlation in breast cancer patients using data from cBioPortal for Cancer Genomics that hosts information from TCGA, (b) Scatter plot of API5 and FGF2 transcript levels showing correlation across different stages of breast cancer i.e., Stage I, Stage II and Stage III (N>133). Quantification was done using the Pearson correlation coefficient (**p<0.01). (c) Scatter plot of API5 and FGF2 transcript levels showing correlation across different molecular subtypes of breast cancer, i.e., Luminal A, Luminal B, HER2-enriched and Basal-like (N>58). Quantification was done using the Pearson correlation coefficient (*p<0.05). (d) Immunoblot analysis showing levels of Api5 and FGF2 across MCF10A isogenic series. (e) Graph showing fold change in Api5, (f) LMW FGF2 (18 kDa) and (g) HMW FGF2 (24 kDa) levels, after normalisation with GAPDH. (h) Agarose gel image showing transcript levels of API5 and FGF2 across MCF10A isogenic series. (i) Graph showing fold change in API5 and (j) FGF2 transcript levels, after normalisation with GAPDH. Statistical analysis was done using One-Way ANOVA (*p<0.05, **p<0.01, ****p<0.0001). Data were pooled from N=3 independent experiments.

To investigate the role of Api5 and FGF2 in the transformation of breast epithelial cells, Api5 and FGF2 protein expression levels were analysed in the MCF10A progression series by immunoblotting. There was a significant increase in Api5 and LMW FGF2 (18 kDa) in MCF10CA1a compared to the MCF10A cells. However, there was no significant difference in Api5 and FGF2 expressions in MCF10AT compared to MCF10A cells (Figure 1d-f). When analysed for HMW FGF2 (24 kDa), no significant difference was observed across MCF10A isogenic series (Figure 1g). Similarly, RT-PCR data revealed elevated transcript levels of API5 and FGF2 in MCF10CA1a compared to MCF10A (Figure 1h-j), concluding that Api5 and FGF2 could be used as reliable indicators for the transformation.

### Api5 regulates FGF2/FGFR1/FRS2α signalling

To study the role of Api5 in regulating FGF2/FGFR1 signalling, the expression of FGF2 was evaluated in Api5-overexpressing MCF10A cells that were grown as spheroid cultures for 16 days and then dissociated [17]. Lysates from these dissociated cells were immunoblotted to confirm Api5 overexpression (Figure S1a). Upon assessing FGF2 in the Api5 overexpression background, a significant increase in LMW FGF2 (18 kDa) and HMW FGF2 (24 kDa) expression was observed (Figure S1a-c). This data was further corroborated by RT-PCR analysis, indicating a significant increase in FGF2 transcript levels upon Api5 overexpression (Figure S1d-f).

Since acinar culture serves as a reliable model for analysing an oncogene’s transformation potential, FGF2 levels were measured upon Api5 deregulation in 3D spheroid cultures. A significant increase in HMW FGF2 (24 kDa) was observed in Api5 overexpressing (hereafter, Api5 OE) spheroid cultures during the early stage, that is, day 4 of morphogenesis (Figure 2a-b). A concomitant increase in the activation of the FGFR1 receptor at tyrosine residue 653/654 was also observed (Figure 2a, c-d). This further led to the phosphorylation of the adaptor protein FRS2α (Figure 2a, e-f), indicating activation of the signalling cascade. Similarly, RT-PCR studies also revealed a significant increase in FGF2 transcript levels upon Api5 OE in 3D day 4 spheroid cultures (Figure 2g-i).

**Figure 2:**
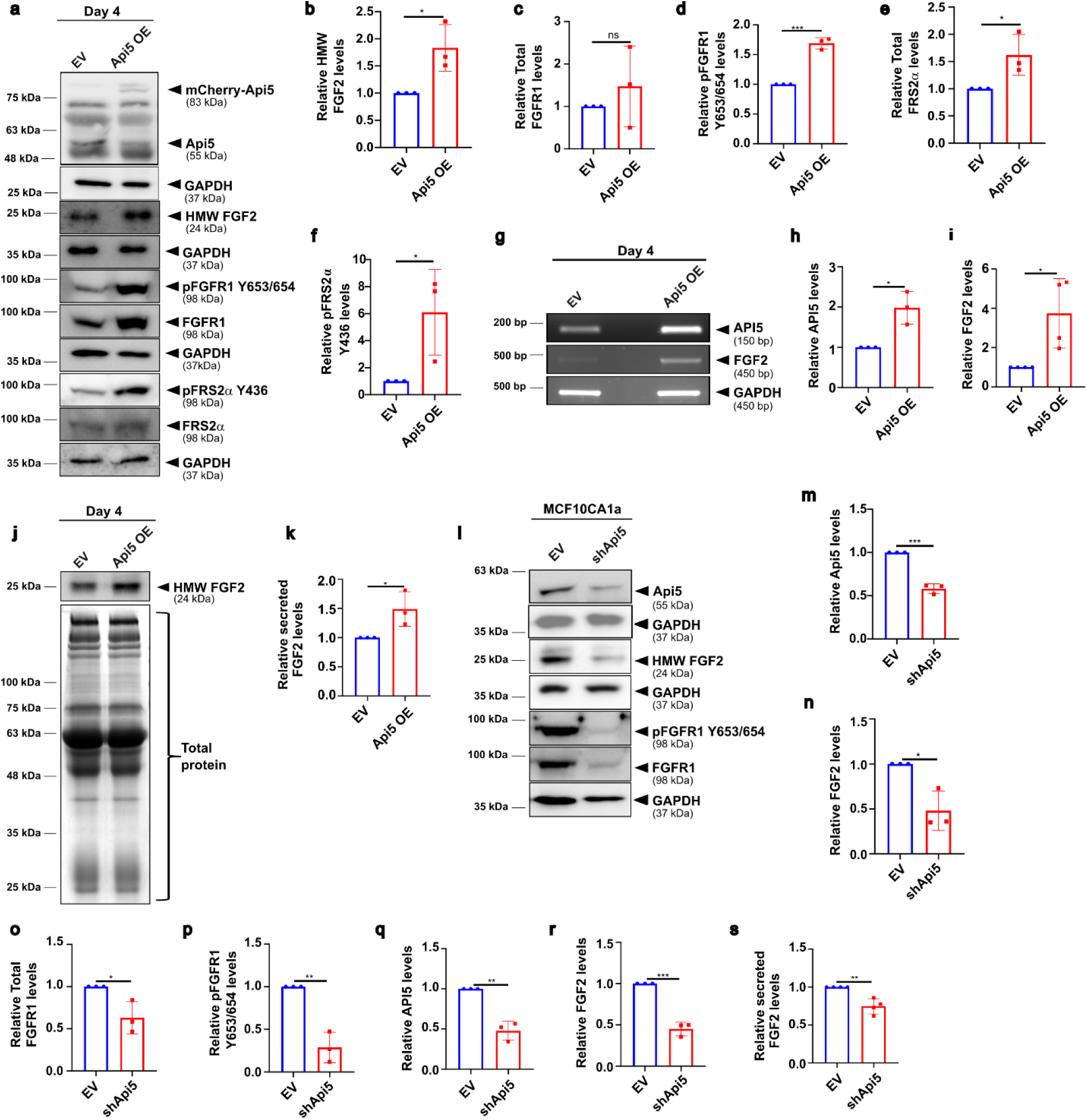
Api5 OE upregulates FGF2/FGFR1/FRS2α signalling axis: (a) MCF10A cells stably expressing mCherry (Empty Vector, EV) and Api5 OE (day 4) in CSII-EF-MCS were lysed, immunoblotted and probed for Api5, FGF2, pFGFR1 Y653/654, total FGFR1, total FRS2, and pFRS2α Y436 levels . Graphs showing relative levels of (b) HMW FGF2 (24 kDa), (c) total FGFR1, (d) pFGFR1 Y653/654, (e) total FRS2α and (f) pFRS2α Y436 upon Api5 OE, normalised to respective totals and GAPDH. (g) Representative agarose gel image showing transcript levels of API5 and FGF2 in day 4 Api5 OE spheroids. Fold change in (h) API5 and (i) FGF2 transcript levels in Api5 OE day 4 spheroids, after normalisation with GAPDH. (j) Immunoblotting performed using conditioned media from Api5 OE 3D day 4 spheroids and probed for FGF2 expression. Total protein in the conditioned media was quantified using Coomassie Brilliant Blue (CBB) staining. (k) Graphs showing relative levels of secreted FGF2 from day 4 Api5 OE conditioned media, normalised with total protein. (l) MCF10CA1a spheroids stably expressing pLKO.1 (Empty Vector, EV) and shApi5 were lysed and immunoblotted and probed for Api5, FGF2, pFGFR1 Y653/654 and total FGFR1-levels. Graphs showing relative levels of (m) Api5, (n) FGF2, (o) total FGFR1 and (p) pFGFR1 Y653/654 in Api5 KD spheroid cultures. Graphs showing fold change in (q) API5 and (r) FGF2 transcript levels in Api5 KD spheroid cultures, normalised to GAPDH. (s) Graphs showing relative levels of secreted FGF2 in Api5 KD conditioned media, normalised to total protein. Statistical analysis was performed using Unpaired Student’s t-test (*p<0.05, **p<0.01, ***p<0.001). Data pooled from N=3 independent experiments.

To assess whether FGF2 was secreted by the cells, immunoblotting was performed to detect its presence in conditioned media from Api5 OE cells grown as spheroid cultures. Increased secretion of HMW FGF2 (24 kDa isoform) was observed from Api5 OE spheroid cultures as early as day 4 (Figure 2j-k). Further, to confirm that overexpression of Api5 increased FGF2 secretion into the media, which then activated the FGFR1/FRS2 signalling axis, conditioned media collected from empty vector and Api5 OE 3D dissociated cells were used to culture MCF10A cells. Cells grown in Api5 OE conditioned media showed increased phosphorylation of FGFR1 (Figure S1g-h), indicating that Api5 OE increases the levels of FGF2, leading to its elevated secretion and activation of FGFR1/FRS2α signalling.

To further confirm that Api5 regulates FGF2/FGFR1/FRS2*α* signalling, Api5 was knocked down (Api5 KD) in the malignant MCF10CA1a breast cancer cells. Downregulation of Api5 was confirmed at the protein and transcript levels (Figure 2l-m, q and S1i). Reduced protein (Figure 2l, n) and transcript levels of HMW FGF2 (24 kDa) (Figure S1i and 2r) were also observed in the Api5 KD spheroid cultures. Additionally, Api5 KD led to reduced secretion of FGF2 in conditioned media (Figure S1j and 2s) and decreased activation of pFGFR1 Y653/654 (Figure 2l, o, p), suggesting that Api5 regulates the FGF2/FGFR1/FRS2α signalling cascade during transformation.

### FGF2/FGFR1 signalling is crucial for Api5 OE-mediated transformation phenotypes of breast epithelial cells

To determine the phenotypic changes associated with increased FGF2/FGFR1/FRS2α signalling in Api5 overexpressed spheroid cultures, the spheroid cultures were treated with 2μM PD173074, a potent FGFR1 inhibitor. Since Api5 was observed to activate FGF2/FGFR1 signalling early, that is, by day 4 of spheroid cultures, PD173074 was added on day 3 for 24 hours (Figure S2a). Reduced activation of pFGFR1 Y653/654 and pFRS2α Y436 was observed at day 12 in PD173074-treated Api5 OE 3D lysates (Figure S2b-d). However, Api5 levels remained unchanged upon addition of the inhibitor, indicating that the inhibition is very specific (Figure S2e-f).

To study the effect of inhibiting the FGF2/FGFR1 signalling cascade on the transformative potential of Api5, morphometric analysis was performed. As previously demonstrated [17], cells overexpressing Api5 formed large spheroids with a fully filled lumen compared with the control acini. A significant increase in volume, surface area, and the number of cells per acini, and a reduction in sphericity were reported. The addition of PD173074 to the Api5 OE spheroids reversed the morphometric changes, resulting in acini similar to those of the control (Figure 3a-e).

**Figure 3:**
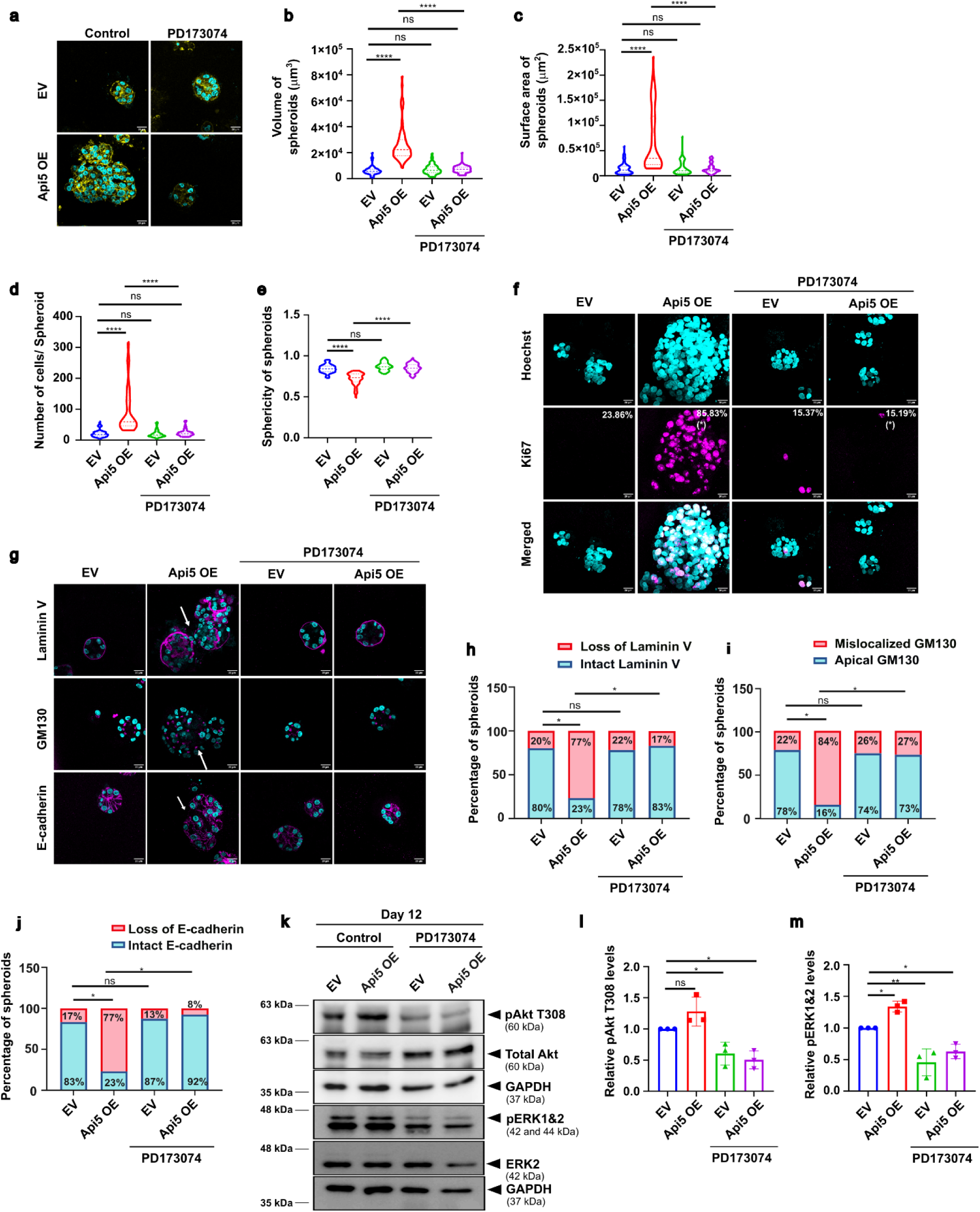
FGF2/FGFR1 inhibition reduces Api5 OE-mediated transformation phenotypes: (a) Representative images of EV (mCherry) and Api5 OE cells grown as 3D “on-top” cultures for 12 days, treated with PD173074 (2μM) and stained with Hoechst 33258 (cyan) as well as Phalloidin (yellow). Graph showing (b) volume, (c) surface area, (d) number of cells/spheroid and (e) sphericity of Api5 OE spheroid cultures upon PD173074 (2μM) treatment. >60 acini from 4 independent experiments were analysed using Huygens Essentials (SVI, Netherlands). Statistical analysis was done using the Kruskal-Wallis test. (f) PD173074 (2μM) -treated day 12 Api5 OE spheroids were fixed and stained with Ki67 (magenta) and Hoechst 33258 (cyan). The percentage of spheroids showing >33% of Ki67-positive nuclei is written in the top right corner of each sample (N=4, n>70). (g) Api5 OE 3D spheroid cultures treated with PD173074 (2μM) and stained with Hoechst 33258 (cyan), Laminin V (magenta), GM130 (magenta) and E-cadherin (magenta). Graphs showing percentage of spheroids with (h) loss of Laminin V, (i) mis-localisation of GM130 and (j) loss of E-cadherin upon PD173074 (2μM) treatment in Api5 OE spheroid cultures (N=4, n>70). Statistical analysis was done using Mann-Whitney test. (k) Immunoblot showing levels of pAkt T308 and pERK1&2 (T185+Y187+T202+Y204) upon PD173074 (2μM) addition in Api5 OE spheroid cultures. Graphs showing fold change in deactivation of (l) pAkt T308 and (m) pERK1&2, normalised to respective totals, which in turn normalised to GAPDH (N=3). Statistical analysis done using One-way ANOVA (*p<0.05, **p<0.01, ****p<0.0001).

To study the effect of FGFR1 inhibition on proliferation, the proliferation marker Ki-67 was used. The Api5 OE spheroids showed increased Ki67 positivity (85.83%) that was reduced to 15.19% upon addition of PD173074 (Figure 3f). Thus, it may be inferred that FGF2/FGFR1/FRS2α signalling may be regulating the altered morphology and enhanced proliferation observed in Api5 OE spheroids. Api5 OE spheroids treated with PD173074 also showed a reduction in the percentage of acini with filled lumen, suggesting a plausible role of FGF2/FGFR1 signalling in apoptosis (Figure S2g, h).

As demonstrated earlier, Api5 OE spheroids were larger and showed disrupted apico-basal polarity and cell-cell junctions compared with control (EV) acini [17]. To determine whether the FGF2/FGFR1 signalling axis regulates this transformed phenotype, the polarity markers were analysed following PD173074 treatment of control and Api5 OE spheroids. 17% of PD173074-treated Api5 OE spheroids showed loss of Laminin V, a basal polarity marker, when compared to Api5 OE spheroids that showed 77% loss of Laminin V (Figure 3g, h). Inhibiting the FGF2/FGFR1 pathway decreased the percentage of spheroids showing mis-localised GM130, an apical polarity marker, from 84% to 27% (Figure 3g, i). Moreover, 77% of Api5 OE spheroids showed loss of the cell-cell junction marker, E-cadherin, which was significantly reduced to 8% upon PD173074 treatment of the Api5 OE spheroids (Figure 3g, j). No change was observed in control cells (EV) after the addition of PD173074 (Figure 3g).

FGF2/FGFR1/FRS2α signalling has been shown to regulate cell survival, proliferation, and apoptosis via various downstream signalling pathways, including PDK1/Akt and Ras/MAPK/ERK [20] [21]. The expression of pAkt T308 and pERK1/2 was studied following PD173074 treatment of Api5 OE spheroids. Activation of both pAkt T308 and pERK1 and 2 (T185+Y187+T202+Y204) was reduced upon inhibiting the FGF2/FGFR1 pathway, indicating the downstream role of the Akt and ERK pathways (Figure 3k-m).

### PDK1/Akt signalling regulates Api5-FGF2-mediated enhanced proliferation and altered morphology during the initial stages of morphogenesis

Various studies have shown that the PDK1/Akt pathway is upregulated in various cancers; thus, levels of pAkt S473 and T308 were initially studied in the MCF10A isogenic series. The Akt pathway was found to be upregulated in the malignant cell line MCF10CA1a compared with MCF10A (Figure S3a-c). Furthermore, the levels of pAkt T308 and S473 were assessed following deregulation of Api5. It was observed that overexpression of Api5 increased the activation of pAkt T308 and S473 in the MCF10A 3D dissociated cells (Figure S3d-f). Moreover, in 3D cultures, Api5 OE in MCF10A increased pAkt T308 levels by day 4 of acinar morphogenesis (Figure S3g, h), while Api5 KD in the malignant cells reduced the levels of pAkt S473 and T308 (Figure S3i-k), indicating that Api5 may play a role in regulating the activation of the PDK1/Akt pathway.

To investigate whether the PDK1/Akt pathway regulated the different morphometric phenotypes, the Api5 OE and control spheroid cultures were treated with 5μM Akt inhibitor on day 3 of morphogenesis for 24 hours. Reduced activation of pAkt T308 was observed on day 12 (Figure S3l-m), with no observable changes in Api5 and FGF2 levels upon Akti addition (Figure 4a-c). Morphometric analysis revealed that inhibition of the Akt pathway led to reductions in volume, surface area, and the number of cells per acini in Api5 OE spheroids (Figure 4d-g). However, Akt inhibition did not have any effect on the number of acini with filled lumen in Api5 OE spheroids (Figure S3n-o). Interestingly, the reduction in the sphericity observed upon Api5 overexpression was also reversed upon Akt pathway inhibition. No changes were observed in the empty vector control upon inhibition of the PDK1/Akt pathway (Figure 4h). Inhibition of the Akt signalling pathway led to only 17.48% Api5 OE spheroids showing Ki67 positivity compared to 80.49% non-inhibited Api5 OE spheroids (Figure 4i). To determine whether Akt signalling plays a role in maintaining cell polarity, Api5 OE and control spheroids were treated with an Akt inhibitor, and the polarity markers were evaluated. Untreated Api5 OE spheroids showed a significant increase in loss of Laminin V (76%), mis-localised Golgi (88%), and loss of E-cadherin (74%) when compared to their respective empty vector controls. However, inhibition of Akt signalling in Api5 OE spheroids had no effect on polarity (Figure 4j-m).

**Figure 4:**
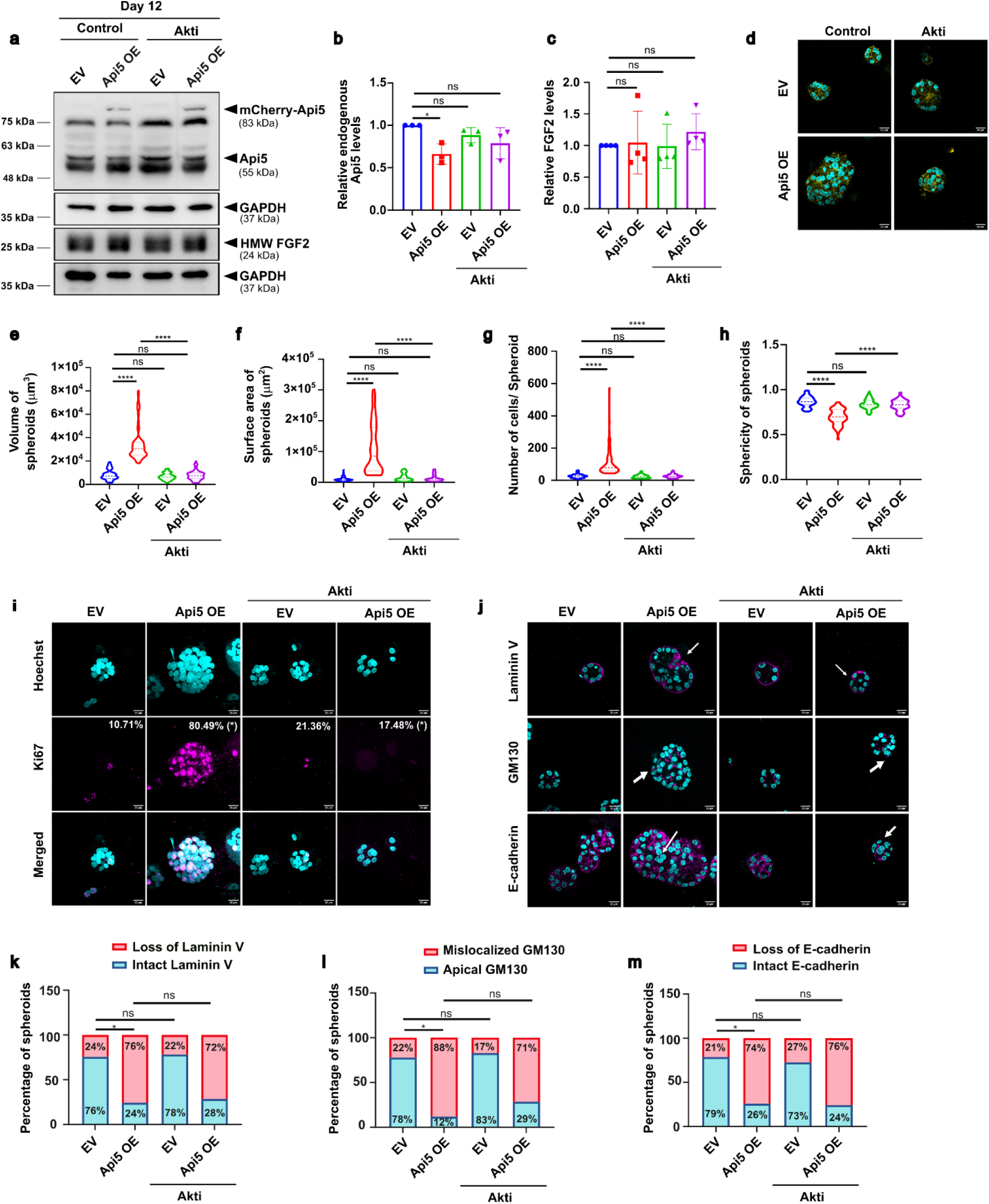
Akt pathway regulates altered morphology and enhanced proliferation via Api5-FGF2 axis: (a) Immunoblot showing protein levels of Api5 and FGF2 in Akti (5μM)-treated lysates of day 12 Api5 OE spheroids. Graph showing fold change in (b) endogenous Api5 (N=3) and (c) FGF2 (N=4); levels normalised to GAPDH. Statistical analysis was done using One-Way ANOVA (d) Representative images of Akti (5μM)-treated empty vector (mCherry) and Api5 OE spheroids grown for 12 days and fixed and stained for Phalloidin (yellow) and Hoechst 33258 (cyan) (Scale bar: 20 μM). Graphs showing (e) volume, (f) surface area, (g) no. of cells/spheroid, (h) sphericity of Api5 OE spheroids that were calculated using Huygens Essentials (SVI, Netherlands). >60 acini from 4 independent experiments were analysed. Statistical analysis was done using the Kruskal-Wallis test. (i) Akti (5μM)-treated Api5 OE day 12 spheroids fixed and stained with Ki67 (magenta) and Hoechst 33258 (cyan). The percentage of spheroids showing >33% of Ki67-positive nuclei is written in the top right corner of each sample (N=4, n>70). (j) Akti (5μM)-treated Api5 OE spheroids stained with Hoechst 33258 (cyan), Laminin V (magenta), GM130 (magenta) and E-cadherin (magenta). Graphs showing percentage of spheroids with (k) loss of Laminin V, (l) mis-localisation of GM130 and (m) loss of E-cadherin upon Akti (5μM)-treatment of Api5 OE MCF10A spheroid cultures (N=4, n>70). Statistical analysis was done using the Mann-Whitney test (*p<0.05, **p<0.01, ***p<0.001 and ****p<0.0001).

### Ras/MAPK/ERK signalling regulates Api5-FGF2-mediated proliferation, lumen filling and polarity disruption during later stages of morphogenesis

First, the expression levels of pERK1&2 (T185+Y187+T202+Y204) were studied upon Api5 deregulation in both non-tumorigenic and tumorigenic cells grown as 3-dimensional cultures. Overexpression of Api5 increased pERK1&2 levels by day 12 (later stages of morphogenesis) in the spheroid cultures compared to the empty vector control (Figure S4a, b), while the knockdown of Api5 led to reduced activation of pERK1&2 (Figure S4c, d). To investigate whether the ERK pathway regulates the transformation of the non-tumorigenic breast epithelial cells with overexpression of Api5, Api5 OE spheroid cultures were treated with 10 μM UO126 on day 11 for the day 12 assays and on days 11, 12, 13, and 14 for day 16 assays (Figure S4e). To confirm ERK pathway inhibition, immunoblotting was performed on lysates collected from day 12 and 16 Api5 OE spheroids treated with UO126. A decrease in pERK1&2 activation was observed upon UO126 addition in both day 12 and 16 Api5 OE spheroids compared to untreated (Figure S4f, g). Moreover, Api5 and FGF2 levels remained unchanged following ERK inhibition (Figure 5a-c).

**Figure 5:**
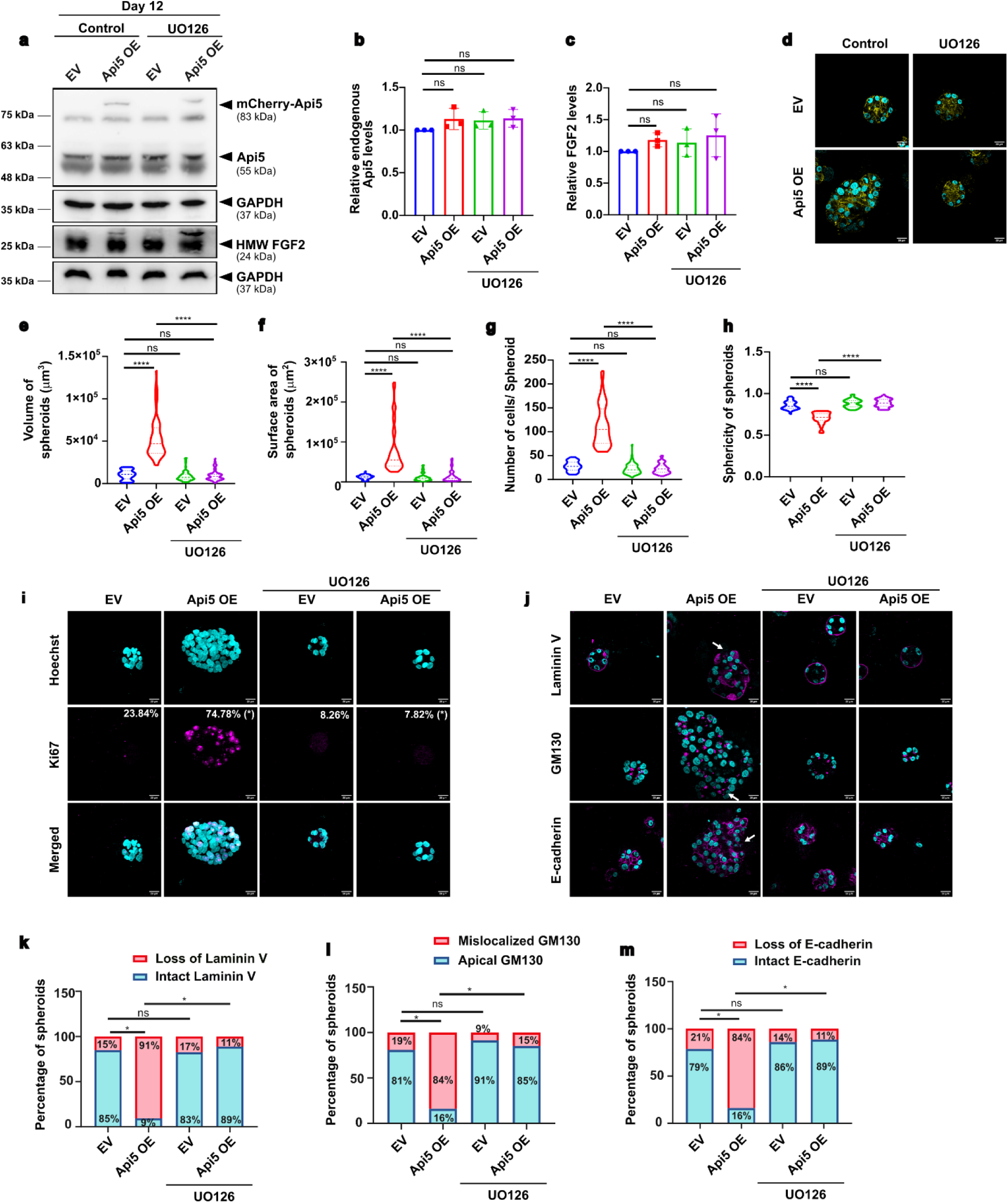
Inhibition of the ERK pathway reduces Api5 OE-mediated transformation of breast epithelial cells during later stages: (a) Immunoblot showing protein levels of Api5 and FGF2 of UO126 (10μM)-treated day 12 lysates from Api5 OE spheroids. Graph showing fold change in (b) endogenous Api5 and (c) FGF2 protein (N=3) levels normalised to GAPDH. Statistical analysis was done using One-Way ANOVA (d). Representative images of UO126 (10μM)-treated empty vector and Api5 OE cells grown as 3D “on-top” cultures for 16 days and fixed and stained for Phalloidin (yellow) and Hoechst 33258 (cyan) (scale bar: 20 μΜ). (e) Volume, (f) surface area, (g) no. of cells/spheroid, (h) sphericity of Api5 OE spheroids were calculated using Huygens Essentials (SVI, Netherlands). >60 acini from 4 independent experiments were analysed. Statistical analysis was done using the Kruskal-Wallis test. (i) UO126 (10μM)-treated day 16 Api5 OE spheroids fixed and stained with Ki67 (magenta) and Hoechst 33258 (cyan). The percentage of spheroids showing >33% of Ki67-positive nuclei is written in the top right corner of each sample. (j) UO126 (10μM)-treated day 16 Api5 OE spheroids stained with Hoechst 33258 (cyan), Laminin V (magenta), GM130 (magenta) and E-cadherin (magenta). Graphs showing percentage of spheroids with (k) loss of Laminin V, (l) mis-localisation of GM130 and (m) loss of E-cadherin upon UO126 (10μM) treatment to Api5 OE spheroid cultures (N=4, n>70). Statistical analysis was done using Mann-Whitney test (*p<0.05, **p<0.01, ***p<0.001 and ****p<0.0001).

**Figure 6:**
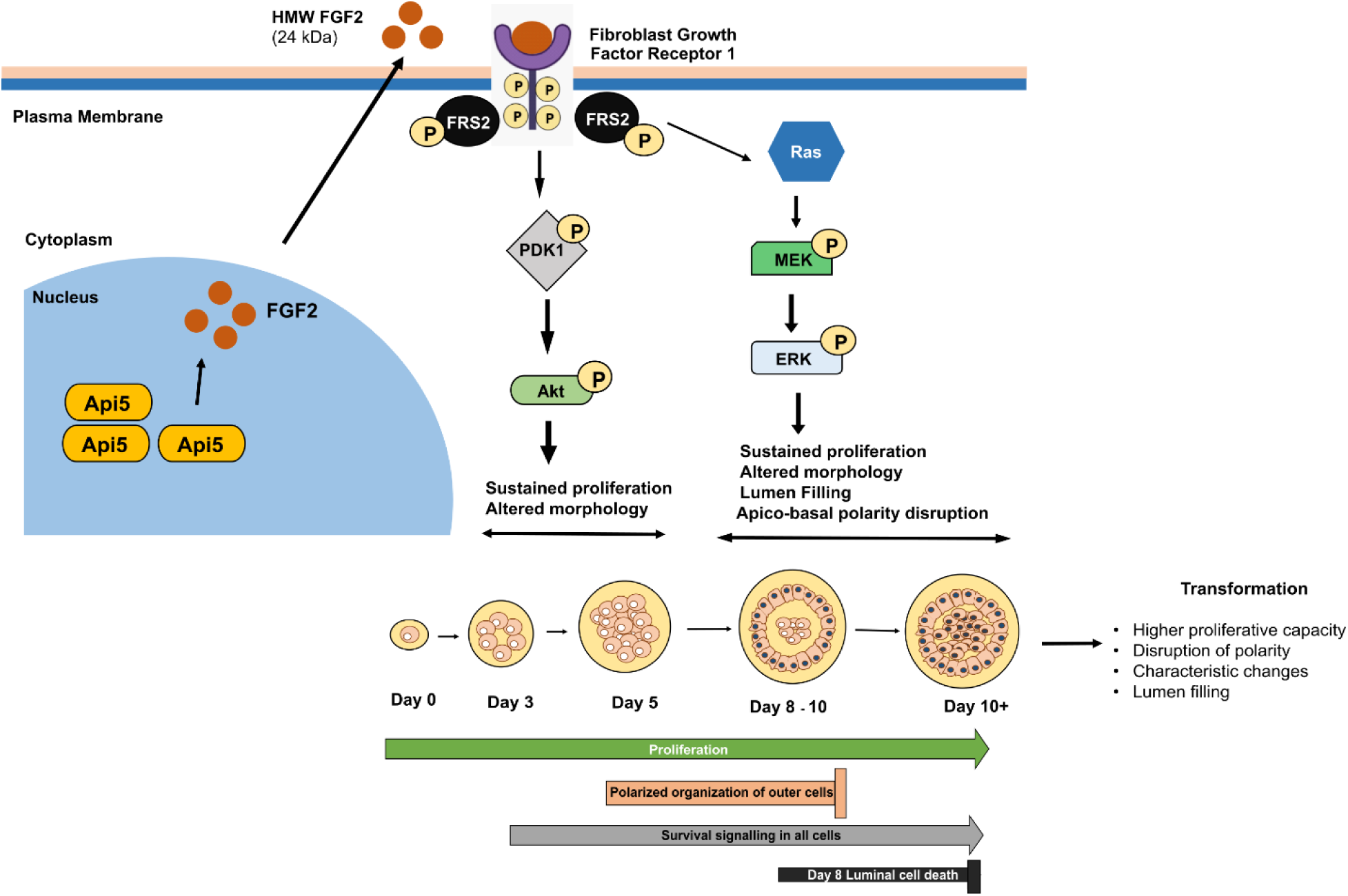
Schematic representation of signalling pathway involved in Api5-FGF2 mediated transformation of breast epithelial cells: Api5 leads to the upregulation of FGF2, which binds to its receptor FGFR1, leading to its autophosphorylation and recruitment of FRS2 adaptor protein. FRS2α, once recruited, gets phosphorylated and activates PDK1/Akt signalling during the initial stages of morphogenesis. This regulates various transformation phenotypes, including altered morphology and enhanced proliferation. Later, this signalling switches to the Ras/MAPK/ERK pathway, regulating hyperproliferation, morphological changes, polarity disruption and lumen formation.

The inhibition of ERK signalling led to reduced volume, surface area, and the number of cells/acini in spheroids overexpressing Api5 (Figure 5d-g). Interestingly, the sphericity is regained in Api5 OE spheroids treated with UO126 (Figure 5h). Api5 overexpression leads to spheroids with filled lumen [17]. However, inhibition of the ERK pathway led to an increase in the percentage of spheroids with hollow lumen in Api5 OE cells (Figure S4h-i). Further, UO126 treatment led to a decrease in cell proliferation in Api5 OE spheroids (7.82%) compared to the untreated spheroids (74.78%) (Figure 5i). Interestingly, UO126 treatment of Api5 OE spheroids during later stages of morphogenesis prevented polarity disruption. In the absence of the ERK inhibitor, 91% and 84% of Api5 OE spheroids showed loss of Laminin V and E-cadherin, respectively, while 84% had mis-localised Golgi. However, treatment of the Api5 OE spheroids with the ERK inhibitor led to a significant decrease in loss of basal (11%), apical (15%) and lateral (11%) polarity (Figure 5j-m).

To check if there was crosstalk between Akt and ERK signalling pathways, pERK1&2 activation upon Akti addition and pAkt T308 levels upon UO126 addition were investigated (Figure S4j-m). No observable changes were detected, thereby indicating the absence of any crosstalk between the two signalling cascades.

## Discussion

Cancer arises from genetic and environmental factors that promote sustained proliferation, evade apoptosis, increase invasion and drive other hallmarks [22]. These gene mutations are often caused by various oncogenes and carcinogens that transform normal cells into cancerous cells. Prior studies have reported Api5 as an oncogenic driver across multiple malignancies. Studies have linked elevated Api5 to poor prognosis and survival outcomes in NSCLC [23] and cervical cancer [24].

Previous studies from our lab have also shown that overexpression of Api5 leads to the transformation of non-tumorigenic breast epithelial cells grown as three-dimensional cultures. Overexpression of Api5 was found to alter morphometry, enhance proliferation, reduce apoptosis, and disrupt the apico-basal polarity of spheroids. These phenotypes were proposed to be regulated by engaging PDK1/Akt early and Ras/MAPK/ERK late in day 16 spheroid cultures [17]. These observations establish Api5’s tumour-promoting potential, setting the stage for unravelling the signalling axis contributing to breast epithelial transformation reported here. *In silico* analysis of patient data performed using the TCGA database revealed that API5 positively correlates with FGF2 in breast cancer. This data was supported by studies of the protein and transcript levels of Api5 and FGF2 in the MCF10A isogenic series. High levels of Api5 and FGF2 are associated with the malignant MCF10CA1a cells as compared to the non-tumorigenic MCF10A cells, indicating that Api5 and FGF2 can serve as biomarkers for breast cancer.

In Matrigel-supported 3D cultures of Api5 overexpressing MCF10A cells, high levels of Api5 upregulated HMW FGF2 at the protein and transcript levels. Interestingly, in addition to increasing FGF2 levels, Api5 was also found to regulate the FGF2/FGFR1/FRS2α signalling axis. Overexpression of Api5 elevated FGF2 secretion by day 4, triggering the phosphorylation of FGFR1 at Y653/654 and the adaptor protein FRS2α at Y436. This was bidirectionally corroborated: conditioned media from Api5-overexpressing spheroids activated FGFR1 in recipient MCF10A cells, while Api5 knockdown in MCF10CA1a spheroids reduced FGF2 expression and secretion, as well as FGFR1/FRS2α phosphorylation. These data refine prior reports of Api5 inducing early FGF2/PDK1-Akt signalling during MCF10A morphogenesis.

Pharmacologic blockade with the FGFR inhibitor PD173074 specifically suppressed pFGFR1/FRS2α without altering Api5 or FGF2 levels. Morphometric aberrations (increased volume, surface area, cells per acinus, reduced sphericity), increased proliferation, lumen filling, and polarity defects were all reverted, alongside reduced activation of downstream pAkt and pERK1/2. This establishes FGF2/FGFR1 as central to Api5’s function in breast epithelial cell transformation.

It is clear that the FGF2/FGFR1 signalling axis is critical for triggering the PDK1/Akt and ERK signalling pathways in Api5-overexpressing transformed cells. Our study has identified the individual pathways associated with the transformation phenotypes observed after overexpression of Api5. Morphometric analysis showed that during the early stages, the Akt pathway regulates the altered morphology of Api5 OE spheroids by restoring their volume, surface area, cell count, and sphericity to levels comparable to those of the control acini. The analysis also revealed that Akt inhibition reduced the proliferative capacity of Api5 OE cells without affecting late-stage polarity or lumen defects. This indicates that Api5-FGF2 drives early growth via the PDK1-Akt pathway. However, by day 12, a signalling switch replaces Akt activity with ERK activation. It is likely that this activated ERK pathway governs the lumen filling and loss of polarity in Api5 OE spheroids. To study the role of the ERK pathway in regulating Api5-FGF2-mediated transformation phenotypes, UO126 (a pERK inhibitor) was used. Our studies reported that blocking the ERK pathway reduced the volume, surface area, and proliferation of Api5-overexpressing cells. MCF10A cells, when grown as acini, acquire apico-basal polarity between days 5 and 8 in 16 days of culture [25]. As previously reported, Api5 overexpression disrupts this polarity [17]. In this study, we demonstrate that the Ras/MAPK/ERK pathway regulates polarity disruption and lumen formation in Api5 OE spheroid cultures. Hence, our data show that Api5 regulates morphology, proliferation, polarity, and lumen filling via the FGF2/ERK pathway during later stages of morphogenesis. The observed Akt-to-ERK switch provides novel mechanistic granularity to Api5-driven transformation. Studies have shown the crosstalk between PDK1/Akt and Ras/MAPK/ERK pathways in cancer cells [26] [27]. Interestingly, in our model system, no crosstalk was observed between the PDK1/Akt and ERK pathways, suggesting that the reversal of transformation phenotypes observed upon inhibition of a particular pathway is largely due to the direct effect of that pathway.

Collectively, these findings position Api5 as a major player channelling FGF2/FGFR1 to regulate PDK1/Akt (early proliferation) and ERK (late polarity loss and lumen filling) without crosstalk. Given the good efficacy of FGFR inhibitors in FGFR-altered breast cancer trials and Api5-FGF2’s prognostic value, targeting this axis in Api5-high/Luminal B subsets via combination therapies requires further investigation.

## Materials and Methods

### Chemicals

Hydrocortisone (H0888), Cholera Toxin (C8052), Epidermal Growth Factor (EGF-E9644), Insulin (I1882), Dimethyl sulphoxide (DMSO, D8418), Triton X-100 (T8787), UO126 monoethanolate (U120-1MG) and PD173074 (P2499) were purchased from Sigma-Aldrich, USA. Phalloidin 488 (A12379) was purchased from Invitrogen (Thermo Fisher Scientific, USA). Matrigel^®^ (356231) and Dispase^®^ (354235) were purchased from Corning, Sigma-Aldrich, USA. Prestained Protein Ladder (MBT092), 50bp DNA Ladder (MBT084-50LN) were purchased from Himedia, India. NaCl (41721), Sodium Lauryl Sulphate (32096) and Tris (71033) were purchased from SRL, India. Paraformaldehyde (AA433689M) and Hoechst 33258 (H3569) were purchased from Thermo Fisher Scientific, USA. Protein Ladder (PLAD03) was purchased from Genes2me, India. AKT1/2 kinase inhibitor (A3153) was purchased from TCI, Japan.

### Cell lines

MCF10A was a generous gift from Professor Raymond C. Stevens (The Scripps Research Institute, California). MCF10AT and MCF10CA1a were purchased from Karmanos Cancer Center, USA.

### Cell culture

MCF10A, MCF10AT and MCF10CA1a were grown as monolayer cultures in growth medium and as 3D cultures in assay medium using standardised protocols [28]. Cells were incubated at 37°C in humidified 5% CO_2_ incubators (Eppendorf, Germany). Dissociated cells were made following the standard protocol [17].

### *In silico* analysis

TCGA (The Cancer Genome Atlas) data was downloaded from Xena Browser (UCSC Xena) by selecting the IlluminaHiSeq dataset along with clinical data. The data merged with expression data based on sample ID using the R package is then sorted to separate data of different molecular subtypes. GraphPad Prism (GraphPad Software, La Jolla, CA, USA) was used to plot the extracted data. Further, the cBioPortal computational database tool was also used for some correlation analysis, using API5 as a "Gene Symbol".

### Immunoblotting

Immunoblotting was performed using specific antibodies according to established protocols [28]. Images were acquired in the ImageQuant LAS 4000 gel documentation system (Cytiva, USA). All immunoblotting data were quantified using a minimum of three independent experiments and have been denoted as fold-difference over respective controls for each blot.

### Immunofluorescence

Immunofluorescence for 3D spheroid cultures was performed using standard protocols [28]. Spheroids were imaged on a Leica SP8 confocal microscope (Leica, Germany) under a 63X oil immersion objective. A minimum of 60 spheroids were captured for morphological characterisation and analysed using Huygens Essential (SVI, Hilversum, Netherlands).

### FGF2 assessment of medium supplements

Api5 OE cells were seeded at a density of 1x10^4^ cells on a Matrigel bed in serum-free growth media and incubated at 37^°^C in a 5% CO_2_ incubator. After 4 days, serum-free conditioned media were collected from both empty vector and Api5 OE 3D spheroid cultures and centrifuged to remove cell debris. This conditioned media was stored at - 80^°^C. On the day of the experiment, the media were kept on ice to facilitate thawing and prevent protein degradation in the conditioned media. The conditioned media was mixed with 6X sample buffer (87 μl of conditioned media + 13 μl of 6X sample buffer) and the required amount i.e., 35 μl in this case was loaded on two 10% SDS-PAGE gels. The gels were run at 120V at room temperature. One SDS-PAGE gel was stained with Coomassie Brilliant Blue, and the other was transferred to a PVDF membrane. The membrane was then probed with an FGF2-specific antibody to assess secreted FGF2 levels in conditioned media and normalised to the total protein load determined by CBB staining. For Api5 KD cells, 7x10^3^ cells were seeded onto a Matrigel bed in serum-free growth media and incubated at 37^°^C in a 5% CO_2_ incubator for 7 days. After 7 days, the conditioned media were collected, and the same procedure was followed.

To study the role of secreted FGF2 in regulating FGFR1 phosphorylation, 0.2×10^6^ Api5 OE 3D dissociated cells were seeded in growth media. After 48 hours, conditioned media were collected and centrifuged to remove cell debris. Simultaneously, MCF10A cells were seeded and grown in growth media. After 24 hours, this growth media was replaced with conditioned media of both empty vector and Api5 OE cells. Cells were allowed to grow for an additional 72 hours, and lysates were collected to assess pFGFR1 Y653/654 levels.

### RNA extraction and cDNA preparation

3D cultures and the MCF10A isogenic series were grown according to standard procedure for RNA extraction [17]. Then, they were harvested by scraping cells following the addition of Trizol (Ambion, USA). RNA extraction was done using the standard protocol. 1μg of RNA was used for the preparation of cDNA using 1ml oligo dT (50mM, Invitrogen, USA) in nuclease-free water, dNTP (2.5mM), and MMLV reverse transcriptase (M0253L, NEB, USA). Further, amplification of the gene of interest was performed using the following PCR conditions: 95^°^C for 60 seconds, 60^°^C (GAPDH, API5), 65^°^C (FGF2) for 45 seconds, 72^°^C for 60 seconds (40 Cycles), and a final extension of 5 minutes. Finally, the amplified product was run on agarose gel (1.5% for API5 and 1% for FGF2). Densitometric analysis was performed using ImageJ software, normalised to GAPDH and then to the control. Primer sequences used are listed in Supplementary Table 3.

### Statistical analysis

At least 3 independent biological replicates were performed for all experiments. Huygens software (SVI, Hilversum, Netherlands) was used to calculate the volume, surface area, number of cells, and sphericity for the 3D spheroid cultures, which were then plotted using GraphPad Prism (GraphPad Software, La Jolla, USA). Student’s t-test or one-way ANOVA was used to analyse the significance of various proteins in Immunoblotting for two or more than two samples, respectively. The Mann-Whitney and Kruskal-Wallis tests were used to assess the significance of morphometric and immunofluorescence analyses.

## Author Contributions

A.G. and M.L. conceived and conceptualised the project. A.G. designed, performed and analysed the experiments. A.G. wrote the draft of the paper, and M.L. edited and wrote the final version of the paper.

## Acknowledgements

We thank Dr. Nagaraj Balasubramanian (IISER Pune, India) and Dr. Rashna Bhandari (CDFD, Hyderabad, India) for their valuable suggestions. The authors acknowledge IISER Pune Microscopy Facility for access to equipment and infrastructure. We also thank the Lahiri Lab members for their useful discussions and comments.

## Supplementary Information

**Supplementary Figure S1:**
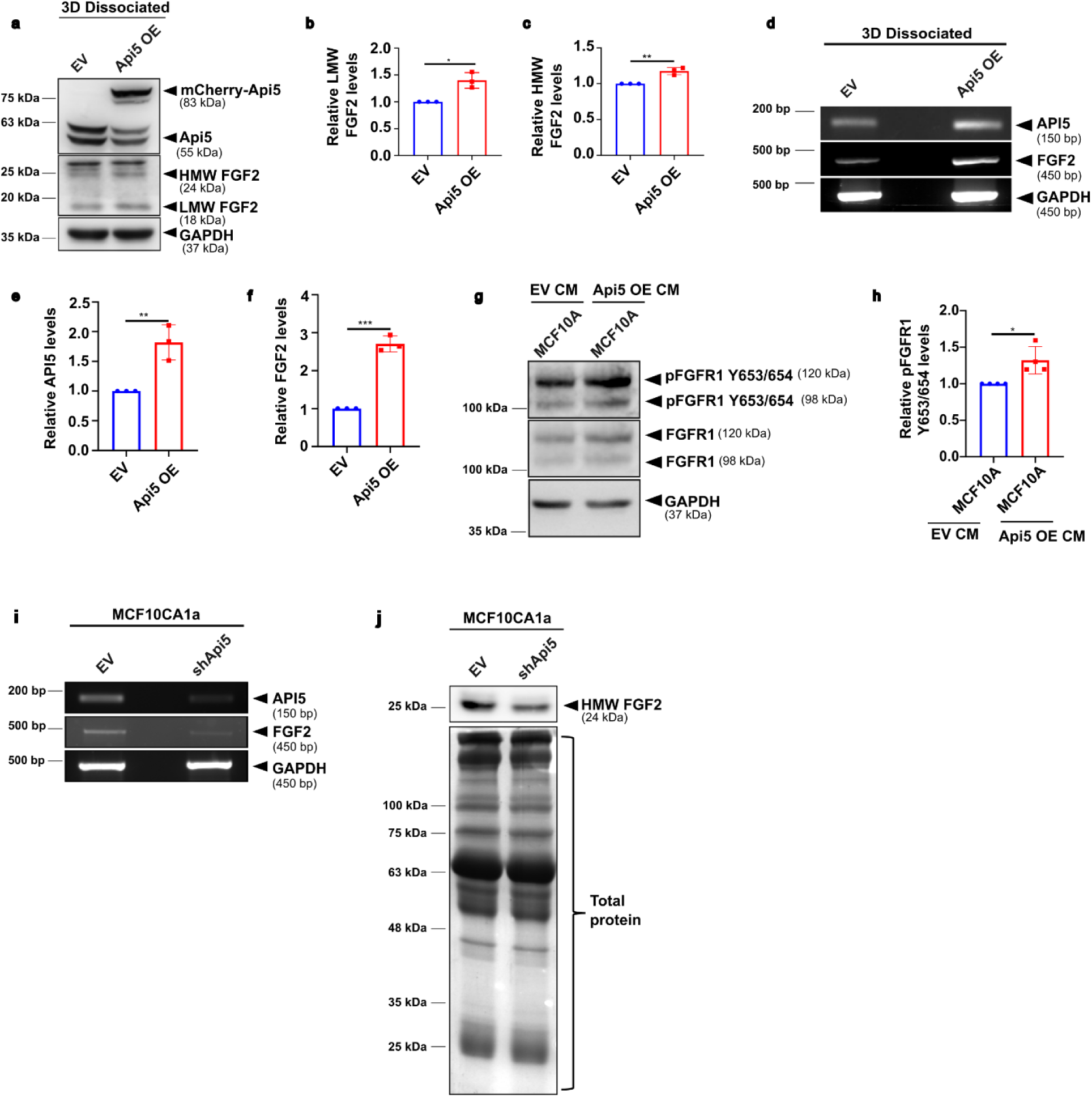
(a) Immunoblot showing Api5 and FGF2 (LMW and HMW) levels upon Api5 OE in 3D dissociated cells. Graph showing relative (b) HMW FGF2 (24 kDa) and (c) LMW FGF2 (18 kDa) levels, after normalisation to GAPDH (N=3). (d) Representative agarose gel image showing transcript levels of API5 and FGF2 in Api5 OE 3D dissociated cells. Graph showing relative levels of (e) API5 and (f) FGF2, after normalisation to GAPDH (N=3). (g) Immunoblot showing levels of pFGFR1 Y563/654 in MCF10A cells grown in conditioned media collected from culturing empty vector and Api5 OE 3D dissociated cells. (h) Graph showing relative levels of pFGFR1 Y653/654 activation in MCF10A grown in Api5 OE conditioned media, normalised to total FGFR1 and GAPDH (N=4). (i) Agarose gel image showing transcript levels of API5 and FGF2 in Api5 KD 3D spheroids. (j) Immunoblot performed with conditioned media from Api5 KD cells and probed for FGF2. Total protein in the conditioned media was quantified using Coomassie Brilliant Blue staining (N=3). Statistical analysis was performed using Unpaired Students t-test (*p<0.05, **p<0.01, ***p<0.001).

**Supplementary Figure S2:**
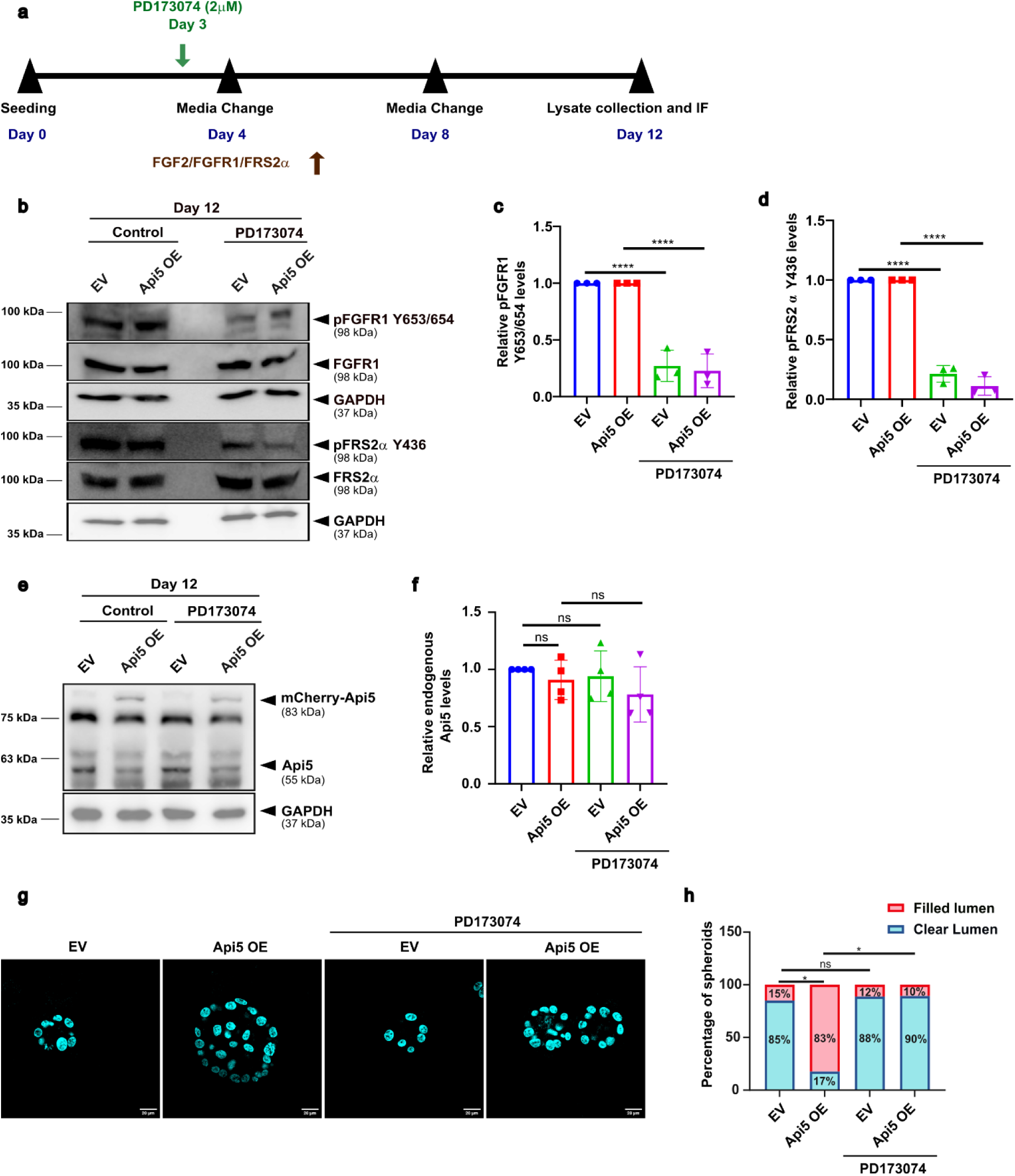
Standardisation of PD173074 in Api5 OE MCF10A 3D cells: (a) Schematic representation of PD173074 (2μM) treatment in 3D spheroid cultures. (b) Lysates collected from PD173074 (2μM) treated Api5 OE, day 12 spheroids and probed for pFGFR1 Y653/654 and pFRS2α Y436, along with corresponding totals and GAPDH. Graph showing relative levels of (c) pFGFR1 Y653/654 and (d) pFRS2α Y436 after normalisation to respective totals and GAPDH (N=3). (e) Immunoblot showing levels of Api5 upon PD173074 (2μM) addition in Api5 OE MCF10A 3D spheroid cultures. (f) Graph showing fold change in endogenous levels of Api5, normalised to GAPDH (N=4). Statistical analysis was done using One-Way ANOVA. (g) Images of PD173074 (2μM) treated Api5 OE 3D spheroids stained with Hoechst 33258 (Cyan) (Central stack) and (h) Graph showing percentage of acini having filled lumen (N=4, n>70). Statistical analysis was done using Mann-Whitney test (*p<0.05, ****p<0.0001).

**Supplementary Figure S3:**
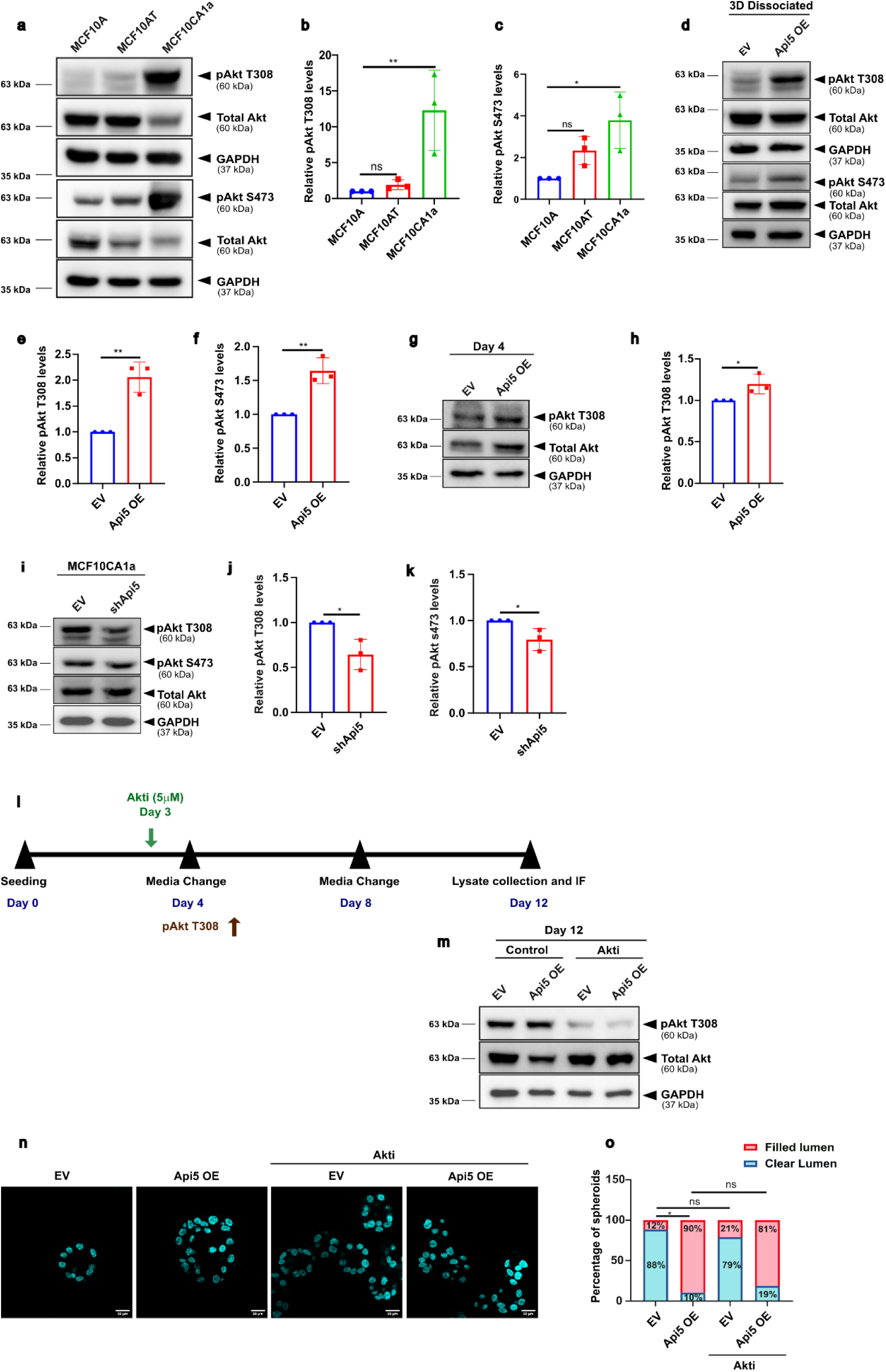
(a) Immunoblot showing pAkt T308 and pAkt S473 levels across MCF10A isogenic series. (b) Graph showing fold change in pAkt T308 and (c) pAkt S473 activation, after normalisation to total Akt and GAPDH (N=3). Statistical analysis was done using One-way ANOVA. (d) Immunoblot showing pAkt T308 and pAkt S473 levels across Api5 OE MCF10A 3D dissociated cells. (e) Graph showing fold change in pAkt T308 and (f) pAkt S473 activation, after normalisation to respective totals and GAPDH (N=3). (g) Immunoblot showing pAkt T308 levels across Api5 OE MCF10A 3D day 4 spheroid cultures. (h) Graph showing fold change in pAkt T308 activation, after normalisation with total Akt and GAPDH (N=3). (i) Immunoblot showing pAkt T308 and S473 levels upon Api5 KD in MCF10CA1a 3D spheroid cultures. (j) Graph showing fold change in pAkt T308 and (k) pAkt S473 deactivation, after normalisation to total Akt and GAPDH (N=3). Statistical analysis was done using Unpaired Students t-test. (l) Schematic representation of Akti (5μM) treatment in Api5 OE 3D spheroid cultures. (m) Lysates collected from Akti (5μM) treated Api5 OE MCF10A 3D D12 cells and probed for pAkt T308, along with corresponding total Akt and GAPDH. (n) Images of Api5 OE 3D spheroids treated with Akti (5μM) and stained with Hoechst 33258 (Cyan) (Central Stack) and (o) Graph showing percentage of acini having filled lumen (N=4, n>70). Statistical analysis was done using Mann-Whitney test (*p<0.05, **p<0.01, ***p<0.001, ****p<0.0001).

**Supplementary Figure S4:**
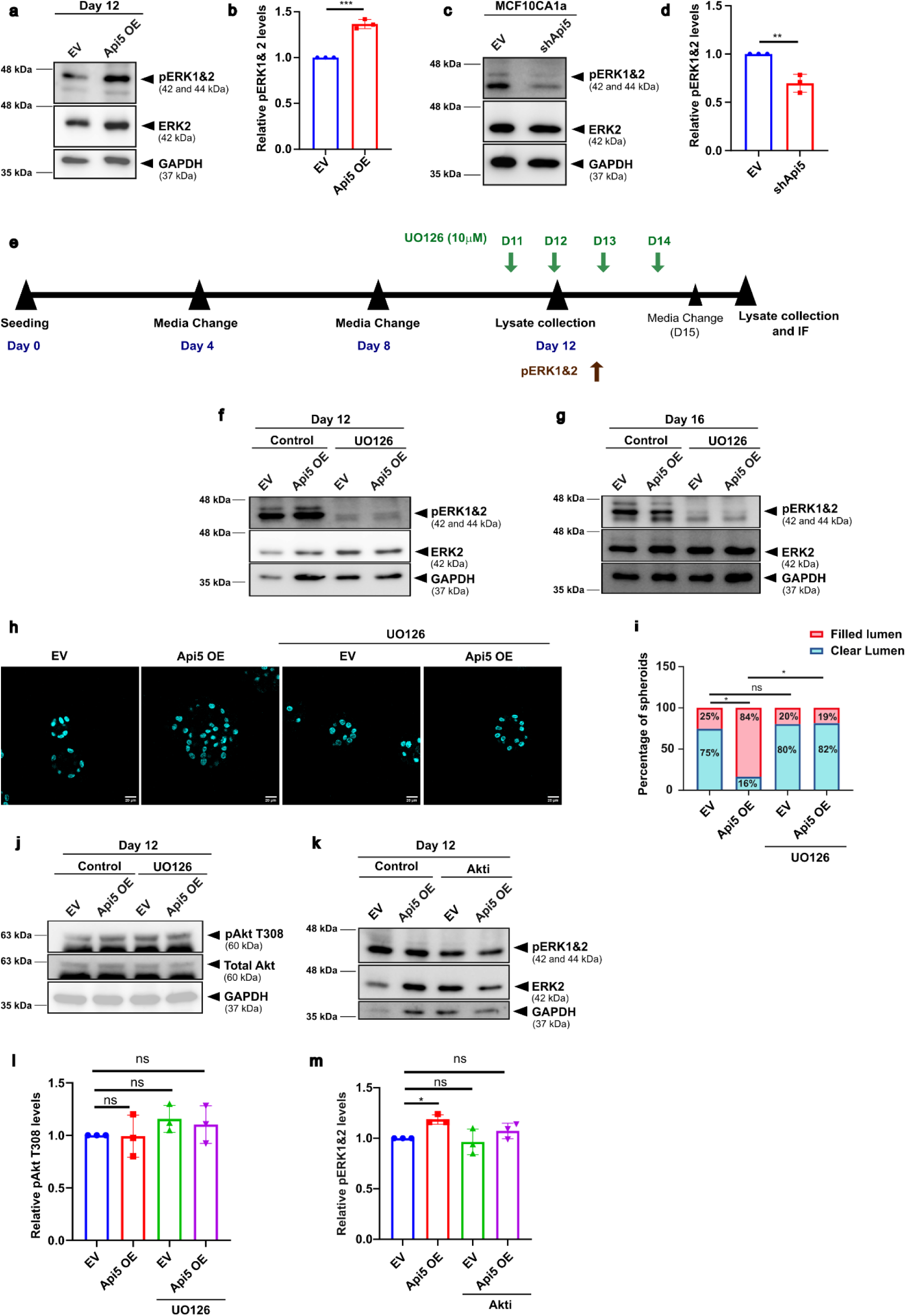
(a) Immunoblot showing protein levels of pERK1&2 in Api5 OE MCF10A 3D day 12 spheroids. (b) Quantification showing fold change in pERK1&2 (T185+Y187+T202+Y204) activation upon Api5 OE, after normalisation with total ERK2 and GAPDH (N=3). (c) Immunoblot showing levels of pERK1&2 upon Api5 KD in MCF10CA1a 3D spheroid cultures. (d) Quantification showing fold change in pERK1&2 (T185+Y187+T202+Y204) deactivation upon Api5 KD, after normalisation with total ERK2 and GAPDH (N=3). Statistical analysis was done using Unpaired Students t-test. (e) Schematic representation of UO126 (10μM) treatment in Api5 OE 3D spheroid cultures for day 12 and day 16. (f) Immunoblot showing levels of pERK1&2 upon UO126 (10μM) treatment in Api5 OE MCF10A 3D spheroids at day 12 and (g) day 16. (h) Images of Api5 OE 3D spheroids treated with UO126 (10μM) and stained with Hoechst 33258 (Cyan) (Central Stack) and (i) Graph showing percentage of acini having filled lumen (N=4, n>70). Statistical analysis was done using Mann-Whitney test. (j) Immunoblot showing levels of pAkt T308 in UO126 (10μM) treated and (k) pERK1&2 levels in Akti (5μM) treated Api5 OE MCF10A 3D day 12 spheroids. (l) Graph showing fold change in pAkt T308 and pERK1&2, after normalisation with respective totals and GAPDH (N=3). Statistical analysis was done using One-way ANOVA (*p<0.05, **p<0.01, ***p<0.001).

**Supplementary Table 1:** Antibodies used for Immunoblotting.

| **Primary antibody** | **Catalogue number** | **Dilution** | **Incubation time** | **Incubation temperature** |
| --- | --- | --- | --- | --- |
| Api5 (Abnova, Taiwan) | PAB7951 | 1:2000 | 3 hours | RT |
| FGF2 (Millipore, USA) | 05-118 | 1:1000 | Overnight | 4°C |
| GAPDH (Sigma Aldrich, USA) | G9545-200UL | 1:10000 (3D)<br>1:40000 (2D, 3D dissociated) | 3 hours (3D)<br>1 hour (2D, 3D dissociated) | RT |
| pFRS2α Y436 (CST, USA) | 3861S | 1:1000 | Overnight | 4°C |
| FRS2α (Santa Cruz, USA) | Sc-17841 | 1:1000 | Overnight | 4°C |
| Phospho-FGF Receptor (Tyr 653/654) (CST, USA) | 3471S | 1:1000 | Overnight | 4°C |
| FGF Receptor 1 (D8E4) (CST, USA) | 9470S | 1:1000 | Overnight | 4°C |
| pAkt T308 (CST, USA) | mAb #4056 | 1:2000 | Overnight | 4°C |
| pAkt S473 (CST, USA) | 9271S | 1:2000 | Overnight | 4°C |
| Akt (CST, USA) | 9272S | 1:2000 | 3 Hours | RT |
| Phospho-p44/42 MAPK (CST, USA) | 4370T | 1:2000 | Overnight | 4°C |
| ERK2 (Abcam, USA) | AB32081 | 1:2000 | Overnight | 4°C |

**Supplementary Table 2:** Primary Antibody used for 3D Immunofluorescence.

| Primary antibody | Catalogue number | Dilution |
| --- | --- | --- |
| Laminin V, anti-Mouse (Merck, USA) | MAB19562 | 1:100 |
| GM130, anti-Mouse (Millipore, USA) | 610822 | 1:100 |
| E-cadherin, anti-Mouse (Abcam, USA) | ab1416 | 1:100 |
| Ki67, anti-Rabbit (Abcam, USA) | ab16667 | 1:100 |

**Supplementary Table 3:**
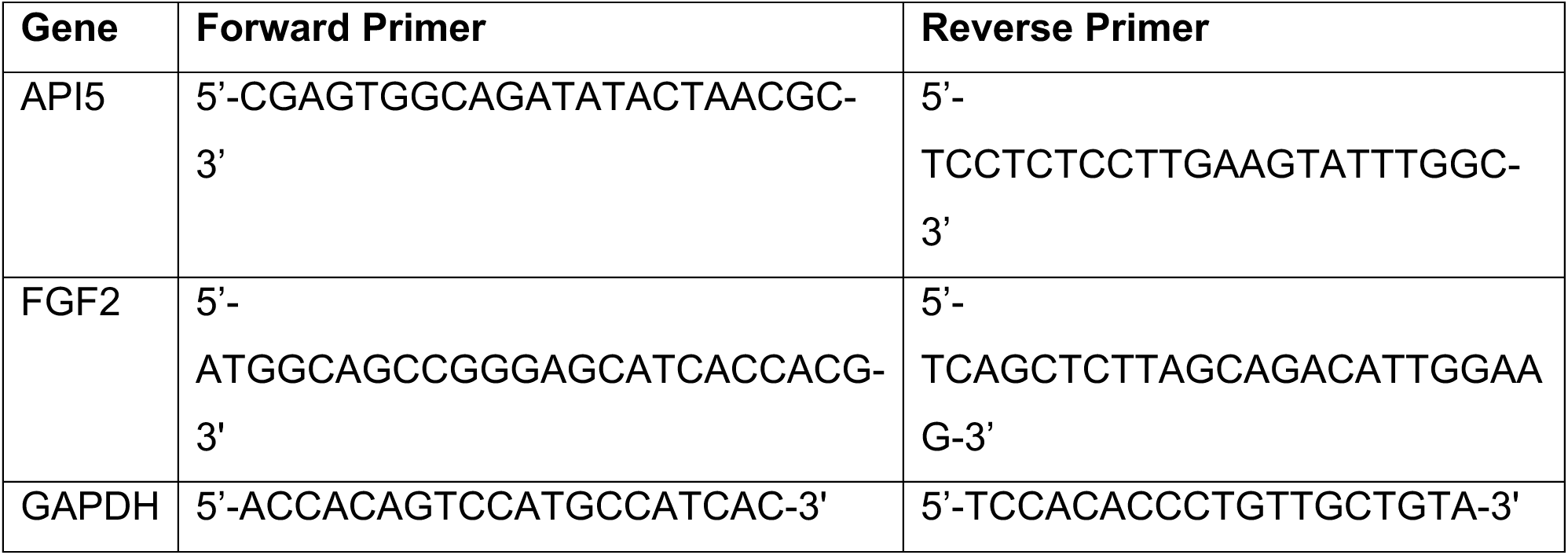
Primer sequences used for RT-PCR.

